# Real-time artificial intelligence prediction of peptide characteristics and MSFragger search improves multiplexed quantification of non-canonical HLA presented peptides in clear cell renal cell carcinoma

**DOI:** 10.64898/2026.05.29.727942

**Authors:** Ana Marcu, Kristin Leskoske, Fengchao Yu, Alexey I. Nesvizhskii, Susan Klaeger, Christopher M. Rose

## Abstract

Non-canonical HLA-presented peptides are promising therapeutic targets, but their low abundance makes them difficult to reproducibly identify and quantify, particularly in multiplexed immunopeptidomics workflows. Here we present MIRA-MS (Model-Informed Real-time Acquisition for Mass Spectrometry), a real-time acquisition strategy that combines fragment ion-indexed database searching with artificial intelligence-based prediction of peptide fragmentation and retention time to guide quantitative scan acquisition. In a clear cell renal cell carcinoma model, MIRA-MS increased the number of quantified non-canonical immunopeptides by 97-107% relative to standard acquisition methods while also improving recovery of canonical peptides by 45-89%. These results establish real-time AI-guided acquisition as a powerful approach for deeper and more reproducible immunopeptidome profiling.

## Main

T cell recognition of peptides presented by human leukocyte antigen (HLA) molecules is a fundamental feature of adaptive immunity, enabling the immune system to detect and eliminate infected or abnormal cells. In cancer, peptides derived from shared or private tumor-specific mutations, known as neoantigens, are important drivers of personalized immunotherapies^1–4^. However, most clinical-stage neoantigen-directed therapies have focused on private mutations in tumors with high mutational burden, with far fewer efforts directed toward tumors with low mutational burden or shared tumor-specific mutations that are infrequently presented across multiple HLA alleles^5^. Interestingly, the observation that some mutation-low tumors, including clear cell renal cell carcinoma (ccRCC), still respond to immune checkpoint blockade (ICB) suggests that additional antigen sources beyond conventional somatic mutations may be immunogenic^6^.

Mass spectrometry–based proteogenomic immunopeptidomics has revealed that tumor cells present not only canonical HLA peptides, but also peptides derived from non-canonical, previously unannotated open reading frames once thought to be untranslated^7–10^. These non-canonical peptides are promising therapeutic targets because they may be shared across patients and can be presented in tumors with low mutational burden^10,11^. Their presentation may also be enhanced by epigenetic perturbation due to modulated transcription and translation of non-canonical sequences^12,13^. Yet their low abundance relative to canonical peptides makes them difficult to reproducibly identify with current immunopeptidomics methods.

To address this analytical challenge, recent methods such as NeoDiscMS have improved label-free detection of immunogenic neoantigens predicted to bind specific HLA alleles, but at the cost of quantifying immunopeptides from other protein sources^14^. Alternatively, isobaric multiplexing improves quantification reproducibility across samples^15^, but it often depends on slower acquisition methods that reduce the overall depth of immunopeptide quantification relative to label-free workflows. In broader proteomics applications, intelligent acquisition methods enabled by instrument application programming interfaces (iAPI), such as inSeqAPI, Orbiter, and MaxQuantLive, have mitigated this tradeoff by allowing real-time peptide identification to dynamically guide instrument acquisition^16–18^ (**Fig. 1a**). Building on this concept, we developed MIRA-MS, a model-informed real-time acquisition strategy that combines AI-based prediction of peptide fragmentation and retention time with a fragment-ion-indexed database search to improve the depth of multiplexed immunopeptidome quantification.

**Figure 1.**
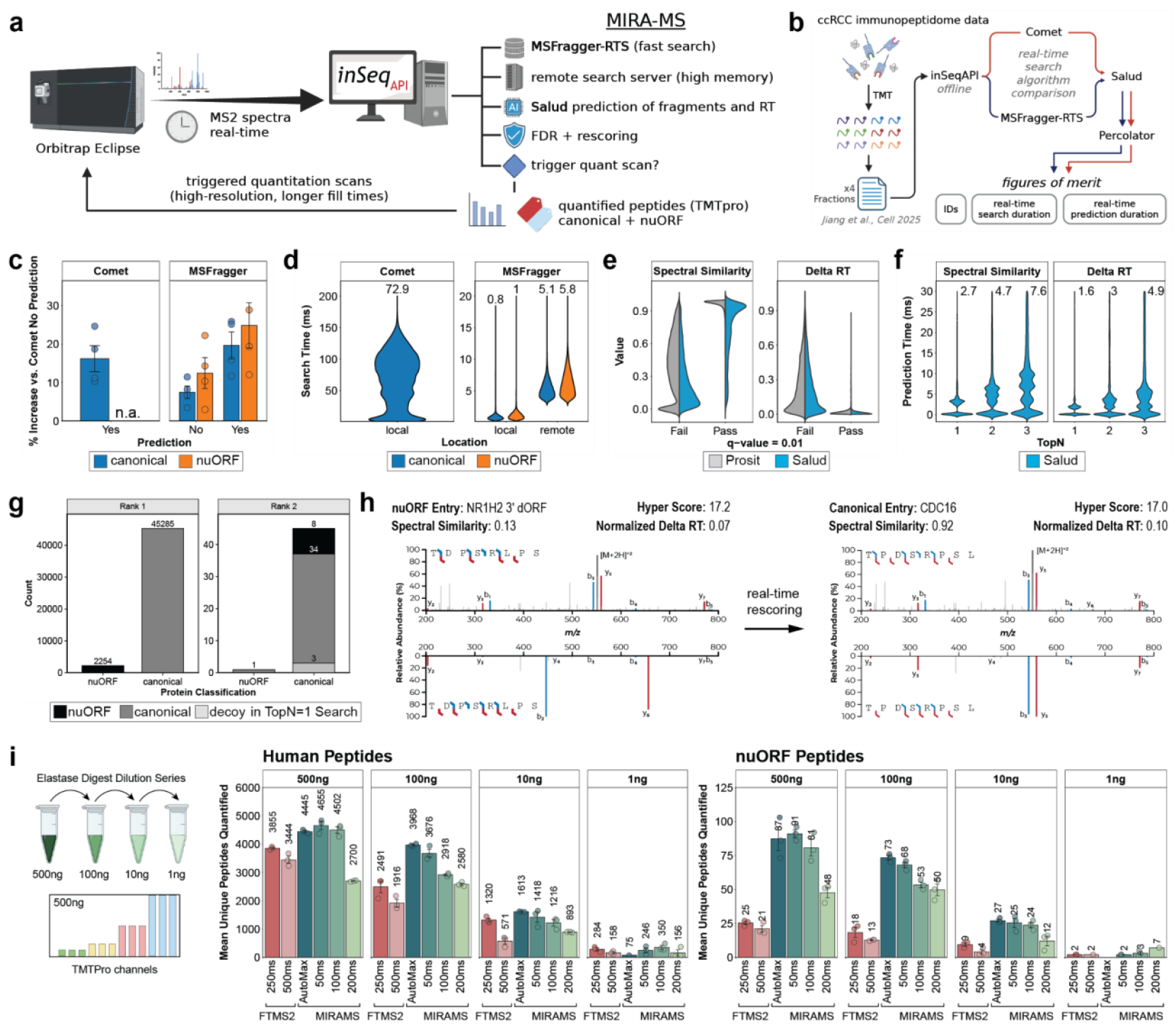
MIRA-MS improves nuORF peptide identification and quantification by combining MSFragger-RTS with Salud real-time prediction. **a)** Overview of MIRA-MS workflow. Instrument directed identification scans are captured by inSeqAPI and analyzed by MIRA-MS through filters including MSFragger-RTS, Salud fragmentation and retention time prediction, and FDR/rescoring before determining if a quantification scan is triggered. **b)** Four fractions from a published ccRCC immunopeptidomics dataset were processed offline through inSeqAPI to benchmark search algorithms computational resource requirements, real-time search and prediction speed, and the number of identified and rescued peptides. Panels (c) - (h) are based on reanalysis of this dataset. **c)** Percentage increase in unique peptide identifications as compared to a Comet search of a canonical database when searching a canonical (blue) or nuORF (orange) database with (‘Yes’) or without (‘No’) using Salud predicted fragmentation and retention time during FDR filtering. Comet search of the nuORF database was not performed as Comet could not create the real-time database file. **d)** Comparison of search times for Comet and MSFragger-RTS (local and remote deployment) when using a canonical (blue) or nuORF (orange) database. Labels indicate median search times (ms). **e)** Comparison of Prosit and Salud prediction values for non-decoy peptide spectral matches. The x-axis denotes peptides that passed or failed 1% FDR filtering. The y-axis ‘Value’ is normalized to the maximum value for each prediction algorithm to enable cross comparison. **f)** Time (ms) required for Salud spectral similarity and retention time predictions when considering one, two, or three peptides per spectrum. Labels indicate median prediction times. **g)** Protein classification (nuORF or canonical) for identified peptides in a search of the nuORF database considering two peptides per spectrum. Colors of stacked bars represent the protein classification for matched spectrum in a search of the nuORF database considering one peptide per spectrum (black = nuORF, dark grey = canonical, light grey = decoy). **h)** Rescoring improves protein classification of peptide identification. A spectrum initially matched to nuORF peptide TDPSRLPS (spectral similarity = 0.13) was reassigned to the correct canonical peptide TPDSRLPS (spectral similarity = 0.92) after Salud rescoring. **i)** Comparison of MIRA-MS and standard peptidomic data acquisition (FTMS2) for serial dilutions of an elastase digest. Unique quantified (cumulative sum S/N ≥ 100) peptides mapping to canonical or nuORF protein entries for standard FTMS2 methods (red) with injection times of 250 ms and 500 ms and MIRA-MS methods (teal) with identification max injection times of 50 ms, 100 ms, 200 ms, and AutoMax. Figure panels a, b, and i were created with biorender.

Most real-time search methods rely on tryptic peptide databases that enable rapid spectral matching (<10 ms), whereas real-time immunopeptide identification requires nonspecific proteome searches with much larger databases and substantially longer search times. To address this challenge, we developed MSFragger-RTS, a real-time implementation of MSFragger that uses fragment-ion indexing to accelerate immunopeptide identification (**Fig. 1a**). Using a previously published multiplexed immunopeptidome dataset^19^, we compared MSFragger-RTS and Comet within inSeqAPI using databases containing either canonical proteins alone or both canonical and non-canonical proteins (nuORF) (**Fig. 1b**). MSFragger-RTS identified more unique peptides than Comet (**Fig. 1c**) and searched the nuORF database with a median time of 1 ms, compared with 72 ms for Comet (**Fig. 1d**). Notably, adding non-canonical sequences had little effect on MSFragger-RTS search time (0.8 ms for canonical versus 1 ms for nuORF), but substantially increased memory requirements, which ranged from 17 to 93 GB depending on the search engine and database used (**Supplementary Fig. 1a**). Despite 128 GB of memory available, Comet could not generate a nonspecific database when nuORF sequences were included, whereas MSFragger required 93 GB of memory to store the indexed fragment ions (**Supplementary Fig. 1a**). Because vendor-supplied instrument computers typically have only 16 GB of memory, local nonspecific real-time searching would require hardware replacement or modification. To overcome this limitation, we configured MSFragger-RTS to operate as a server on a remote high-memory computer. Remote analysis introduced only a 5 ms delay, enabling nonspecific nuORF database searches in 6 ms overall (**Fig. 1d**).

Although this architecture enabled sufficiently rapid real-time searching, accurate immunopeptide identification has been further improved by incorporating predictions of peptide fragmentation and retention time in post-acquisition false discovery rate filtering. To bring these benefits into real-time acquisition, we developed Salud, a Prosit-derived model adapted for ONNX conversion and native deployment within inSeqAPI. We benchmarked Salud against Prosit using spectral similarity and delta RT across all non-decoy identifications and observed comparable performance between the two models (**Fig. 1e**). Encouragingly, incorporating Salud prediction scores increased unique immunopeptide identifications by an average of 10% for MSFragger-RTS searches of both canonical and nuORF databases (**Fig. 1c**).

Search engines often report more than one candidate peptide for each spectrum (TopN), and AI-based rescoring using predicted fragmentation and retention time can promote a lower-ranked match to the top hit. Because Salud improved overall immunopeptide identification, we next asked whether it could support real-time rescoring of multiple candidate peptide matches per spectrum. This capability is particularly important for nuORF-inclusive databases, where the larger search space increases the risk of false-positive nuORF assignments. To enable rescoring, fragmentation and retention time must be predicted for each candidate peptide associated with a spectrum. Encouragingly, local de novo fragmentation prediction with Salud was sufficiently fast for real-time application, with spectral similarity rescoring requiring 2.7-7.6 ms and delta RT requiring 1.6-4.9 ms for 1-3 candidate peptides, respectively (**Fig. 1f**). To further accelerate analysis, inSeqAPI cached prior Salud predictions, reducing prediction time to 0.2 ms for previously observed peptides (**Supplementary Fig. 1b**). Because third-ranked peptides did not pass false discovery thresholds (**Supplementary Fig. 1c**), subsequent analyses considered only the top two candidates per spectrum. Using MSFragger-RTS, we found that rescoring promoted 46 second-ranked peptides to the top position, and 17% (8/46) of these were reassigned from nuORF to canonical sequences (**Fig. 1g**). For example, a spectrum initially assigned to the nuORF peptide TDPSRLPS with low spectral similarity (0.13) was reassigned in real time to the second-ranked canonical peptide TPDSRLPS, which showed substantially higher spectral similarity (0.92) (**Fig. 1h**). Together, these results show that real-time rescoring is fast enough for online use and improves canonical-versus-nuORF assignment.

We next integrated a fragment ion-indexed search (MSFragger-RTS) with real-time Salud-based spectral similarity and delta RT prediction to create MIRA-MS (Model-Informed Real-time Acquisition for Mass Spectrometry). For each identification scan received by inSeqAPI, MIRA-MS performs an MSFragger-RTS database search, Salud-based prediction, and FDR filtering/rescoring before triggering quantitative scans (**Supplementary Fig. 2a,b**). After optimizing key parameters, including real-time FDR, protein priority, maximum injection time, real-time prediction, identification scan resolution, and offline search settings (**Supplementary Fig. 3a-f**), we applied MIRA-MS to a TMTpro-labeled unspecific digest dilution series to model HLA-like peptides. Relative to standard FTMS2, MIRA-MS increased the number of quantified canonical peptides by 20-60% and quantified nuORF peptides by 200-300% (**Fig. 1i; Supplementary Fig. 4**). MIRA-MS improvements reflect the use of rapid, lower-resolution identification scans to prioritize precursors for longer, higher-resolution quantitative scans. Although the primary purpose of the quantitative scans is reporter ion measurement, they also provide higher-quality MS/MS spectra with greater ion accumulation, resulting in improved MSFragger hyperscores (**Supplementary Fig. 5**).

Having established that MIRA-MS improves multiplexed quantification in a controlled benchmark setting, we next asked whether it could enhance detection of shared HLA-presented peptides in a biologically relevant tumor model. To test this, we isolated HLA-I peptides from three ccRCC cell lines treated with DMSO or the HDAC inhibitor Vorinostat (**Fig. 2a, Supplementary Table 2**). We first enriched HLA-A*02:01-specific peptides from ∼10e6 cells to increase recovery of peptides sharing the same binding motif from one of the most frequent alleles, labeled the eluted peptides with 18-plex TMTpro, fractionated them, and analyzed them using either standard FTMS2 or our optimized MIRA-MS method (**Fig. 2a**). Relative to the control method, MIRA-MS quantified 1,422 additional canonical peptides (45% increase) and 63 additional non-canonical peptides (97% increase) (**Fig. 2b**). Among all peptides identified by MIRA-MS, 2.7% were nuORF-derived, consistent with previous studies of non-canonical immunopeptidomes. These nuORF peptides spanned multiple classes of non-canonical transcripts, with most annotated as lncRNA-derived or originating from annotated ORFs absent from the UniProt reference proteome (**Fig. 2c**). Both canonical and nuORF peptides quantified by MIRA-MS matched the expected A*02:01 binding motif, and most peptides in both groups were predicted binders (**Fig. 2d,e**).

**Figure 2:**
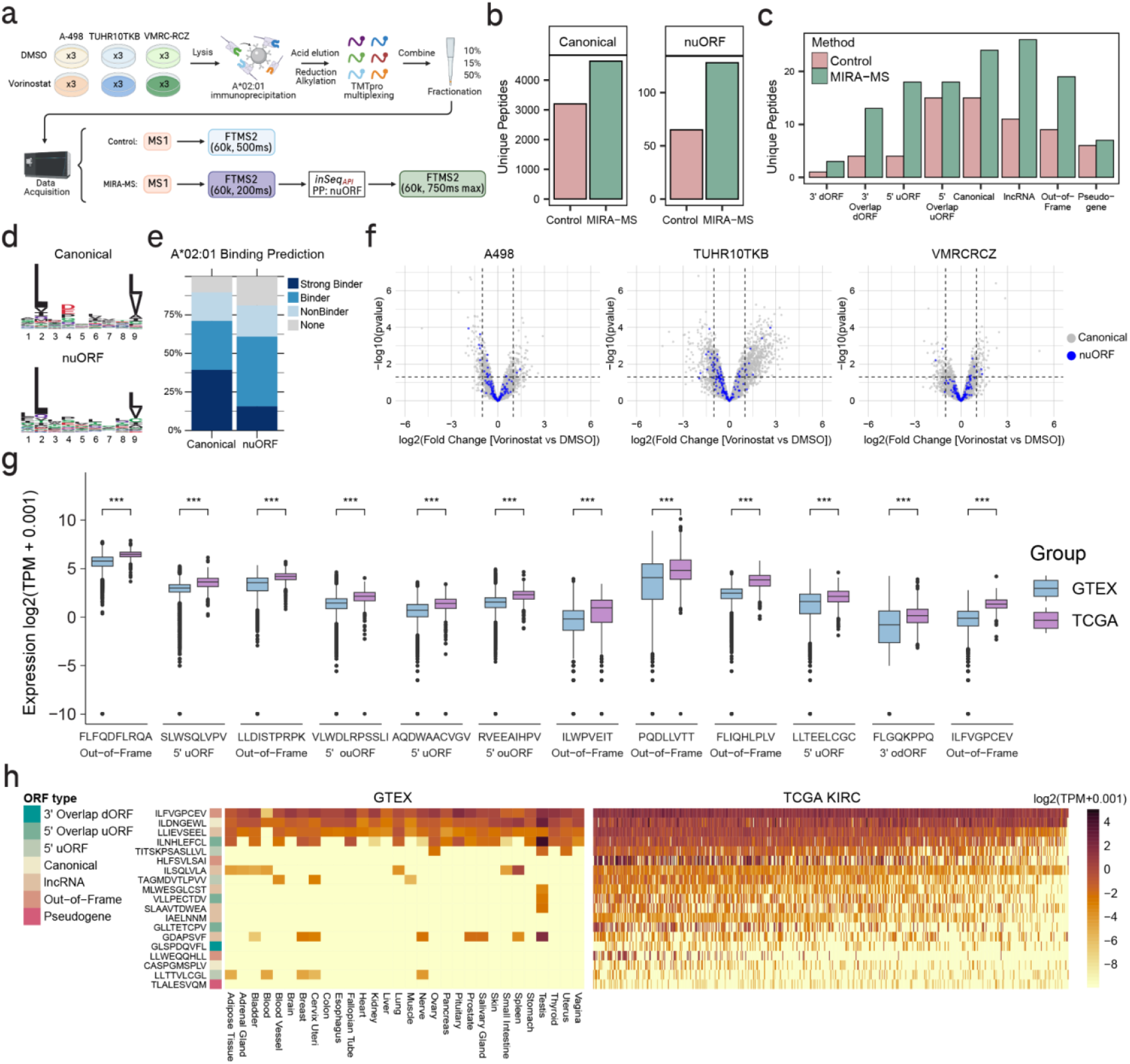
HDAC inhibitor treatment alters HLA-peptide abundance in clear cell renal cell carcinoma. a) ccRCC cell lines A-498, TUHR10TKB and VMRC-RCZ were treated, in triplicate, with either 5 µM Vorinostat or DMSO for 18 hr. A*02:01 HLA-I peptide complexes were immunoprecipitated with BB7.2 antibody, acid eluted, multiplexed with TMTpro, fractionated and analyzed on an Orbitrap Eclipse mass spectrometer using either control FTMS2 or optimized MIRA-MS acquisition. Figure created with bioRender.(b) Number of unique quantitated peptides mapping to canonical or nuORF databases for each method. (c) Number of unique nuORF peptides by type and acquisition method. (d) Sequence logos and (e) predicted A*02:01 binding of canonical and nuORF peptides quantitated using MIRA-MS. (f) Differential abundance of canonical (grey) and nuORF (blue) peptides in Vorinostat vs DMSO treated cell lines. (g) Expression levels of tumor-associated transcripts in TCGA KIRC vs GTEx normal tissue. ***p < 0.05 by Wilcox test. (h) Expression levels of tumor-specific transcripts mapping to identified A*02:01 peptides.

Beyond the A*02:01-enriched analysis, MIRA-MS also improved peptide identification in pan-HLA-I-enriched samples, quantifying 1,640 additional canonical peptides (89% increase) and 56 additional nuORF peptides (107% increase) relative to standard FTMS2 analysis (**Supplementary Fig. 6a**). Peptide abundance varied across cell lines and correlated with predicted HLA-I allele binding, underscoring the importance of shared HLA alleles across samples for multiplexed experiments (**Supplementary Fig. 6b**). Vorinostat treatment induced global proteome changes (**Supplementary Fig. 7**) and altered the abundance of 60 A*02:01-restricted nuORF peptides (**Fig. 2f**). Of these, 12 were encoded by transcripts with higher median expression in kidney tumors than in normal tissues (**Fig. 2g**). In addition, regardless of treatment, 19 A*02:01-restricted nuORF peptides identified with MIRA-MS were encoded by tumor-specific transcripts (**Fig. 2h**), though the actual biological or immunogenic effect of these nuORF peptides remains unknown.

In conclusion, MIRA-MS establishes that real-time AI-guided prediction of peptide fragmentation and retention time can improve peptide assignment and deepen multiplexed immunopeptidome quantification across samples and treatments, with particular benefit for low-abundance non-canonical HLA-presented peptides. Beyond immunopeptidomics, this framework may enable broader applications of intelligent data acquisition, including improved quantification of post-translational modifications and automation of highly multiplexed targeted proteomics.

## Methods

### MSFragger-RTS

We extended the fast database search engine MSFragger to support real-time searching and designated this implementation MSFragger-RTS. Because instrument control workstations typically have limited CPU and memory resources, we implemented a client– server architecture in which inSeqAPI functions as the client and MSFragger-RTS runs as the server. This design supports deployment either on the instrument-attached computer (local mode) or on a separate network-accessible high-performance computer (remote mode). During data acquisition, inSeqAPI streams each MS/MS scan to MSFragger-RTS and synchronously receives the search result. Upon receiving a scan, MSFragger-RTS performs spectrum preprocessing and database searching using the same core codebase as MSFragger, ensuring algorithmic consistency between offline and real-time workflows. To minimize communication latency between inSeqAPI and MSFragger-RTS, we adopted gRPC (https://grpc.io/), a high-performance remote procedure call framework with multi-language support, including Java and C#. A demonstration of client–server communication is available at https://github.com/Nesvilab/MSFragger-RTS-client.

### Salud, a Prosit-derived models for MS/MS fragmentation and iRT prediction

Salud comprises two deep learning models that were implemented in TensorFlow/Keras and adapted from the Prosit framework^20,21^ to predict (i) peptide fragment ion intensities and (ii) normalized retention times (iRT). Peptide sequences were represented as fixed-length integer-encoded vectors (length = 30; padded/truncated as needed) and embedded into a 32-dimensional latent space using a trainable embedding layer (vocabulary size = 30).

### Salud Fragmentation model

Fragment ion intensity prediction was performed using an encoder–decoder recurrent neural network conditioned on precursor/acquisition metadata. The peptide sequence embedding was processed by a bidirectional GRU encoder (256 units per direction, return sequences), followed by dropout (rate 0.1), and a second bidirectional GRU (256 units per direction, return final state), followed by dropout (rate 0.1), yielding a 512-dimensional peptide representation.

To condition predictions on acquisition settings, three auxiliary inputs were provided: precursor charge state (6-dimensional vector), normalized collision energy (NCE) (scalar), and collision mode (2-dimensional indicator). These inputs were concatenated and projected through a linear dense layer (512 units). The resulting 512-dimensional conditioning vector was combined with the peptide representation via element-wise multiplication to yield a context-conditioned latent representation.

The conditioned latent vector was repeated across 29 positions (corresponding to the maximum number of backbone cleavage sites for length-30 peptides) and passed through a bidirectional GRU decoder (256 units per direction, return sequences), followed by dropout (rate 0.1). A time-distributed dense layer produced 6 outputs per position, followed by a LeakyReLU activation; outputs were flattened to form the final predicted fragment intensity vector.

The model was trained using Adam with AMSGrad (learning rate 1×10^−3^) and a masked spectral distance loss to compare predicted and observed spectra while appropriately handling invalid/padded fragments. In contrast to Prosit’s original sequence-to-sequence formulation, the attention mechanism was omitted; conditioning was implemented through multiplicative interaction between the encoded peptide representation and the acquisition-parameter embedding. The model and post-training weights were converted to a ‘.onnx’ format using tf2onnx (v1.16.1) and natively integrated into inSeqAPI using the OnnxRuntime nuGet package (v1.21).

### Salud iRT model

A separate recurrent neural network was used to predict iRT from peptide sequence alone. The embedded sequence was processed by a bidirectional GRU (256 units per direction, return sequences) followed by dropout (rate 0.5), then summarized by a unidirectional GRU (512 units, return final state) with dropout (rate 0.5). A dense regression head (dense 512 units with ReLU, followed by LeakyReLU and dropout rate 0.3) mapped the latent representation to a single scalar iRT prediction via a final linear dense layer. The model was trained using mean squared error loss and Adam with AMSGrad (learning rate 1×10^−3^). The model and post-training weights were converted to a ‘.onnx’ format using tf2onnx (v1.16.1) and natively integrated into inSeqAPI using the OnnxRuntime nuGet package (v1.21).

### Analysis of previous multiplexed immunopeptidomics data

TMT labeled ccRCC immunopeptidomics data from Jiang et al. were downloaded from MassIVE under the identifier MSV000096406^19^. Four fractions of TMT labeled immunopeptides were analyzed using inSeqAPI in offline mode. For the searches we used either the single search function of Comet (v2025.0.20) natively integrated into inSeqAPI or MSFragger-RTS operated locally on a high-performance PC (128 GB memory) or remotely on a Windows server (512 GB memory). For analyses that utilized Salud prediction, the PredictSpectraFilter and PredictRTFilter were implemented before the FDRFilter. The FDRFilter of inSeqAPI outputs a Percolator input file at the end of execution and the resulting peptide spectral matches were filtered for false discovery rate of 1% within Percolator (v3.5). The resulting identifications were further filtered for unique peptide sequences for comparison of algorithm performance. A custom R script was used to extract execution times of Salud spectral similarity and delta RT calculations from inSeqAPI’s ‘offline_analysis.csv’ files.

To compare the performance of Salud and Prosit, raw files were also searched within FragPipe with identical search parameters and utilization of Prosit (Prosit_2020_intensity_TMT, Prosit_2020_irt_TMT) within MSBooster. The resulting percolator input file from FragPipe was used to extract the unweighted spectral entropy and delta RT loess real columns. The delta RT loess real values were normalized to the largest value resulting in a range from 0 to 1. These values were compared to the spectral similarity and delta RT (also normalized to the largest value) values from Salud in cases where the raw data file name, scan number, and peptide sequence matched. For plotting the values were separated into peptides below and above a q value of 0.01 following Percolator (v3.5) analysis.

### Elastase Digest Dilution Series

To model the nonspecific termini characteristic of HLA-presented peptides, an elastase digest was generated from 1e8 C1R HLA-A*02:01 monoallelic cells. Cells were lysed in 1 ml lysis buffer (8M urea, 50 mM Tris pH 9.0) followed by brief sonication. The lysate was clarified by centrifugation (20,000g, 10 minutes, 10°C). A 2 mg aliquot of the protein lysate was reduced with 5 mM Dithiothreitol (DTT, Thermofisher, cat. A39255) for 30 minutes at RT and alkylated with 15 mM iodoacetamide (IAA, Sigma-Aldrich, cat. I1149-5G) for 45 minutes in the dark at RT. The sample was diluted to 1 M urea using 50 mM Tris pH 9.0 and digested with 40 µg elastase (Promega, cat. V189A) for 1h at 37°C on a shaker at 500 rpm. The resulting peptides were acidified with TFA and desalted using Sep-Pak (200 mg tC18; Waters).

Peptides were vacuum-dried and resuspended with 100 mM HEPES to a concentration of 1.3 mg/ml. The TMTPro 18-plex (Thermofisher, cat. A52045, lot: ZD386952) reagent was reconstituted with 10 µl anhydrous acetonitrile (Sigma-Aldrich, cat. 900644-4×2ml). Defined peptide amounts (100 µg, 50 µg, 20 µg and 10 µg) were labeled in triplicates for 1h at RT, 500 rpm. Following quenching with 5% hydroxylamine, labeled peptides were combined, dried, and desalted using Sep-Pak (50 mg C18 SepPak; Waters). Labeled peptide mixtures were dried and stored at −80°C prior to analysis.

To monitor the performance of our methods at different sample inputs, we serially diluted the TMTPro-labeled peptide mix to concentrations of 500 ng/µl, 100 ng/µl, 10 ng/µl and 1 ng/µl.

### Liquid chromatography-tandem mass spectrometry (LC-MS) analysis

Peptides were reconstituted in buffer A (2% acetonitrile/0.1% formic acid) and separated using a Dionex Ultimate 3000 RSLCnano system (Thermo Fisher Scientific, Inc.) coupled to an Orbitrap Eclipse Tribrid mass spectrometer (Thermo Fisher Scientific, Inc.). Chromatographic separation was performed on an Aurora XT column (Ion Opticks, 25 cm × 75 μm, 1.7 μm C18) at a flow rate of 300 nl/min. Peptides were eluted over a 45 min linear gradient from 4% to 30% buffer B (98% acetonitrile/0.1% formic acid).

### Instrument Methods (Control FTMS2, Control FTMS2-MS3, Control ITMS2-FTMS3)

All methods employed FAIMS with compensation voltages at -40 V and -60 V. MS1 scans were acquired in the Orbitrap at 120,000 resolution across a mass range of 350–1,350 m/z with a normalized automatic gain control (AGC) target of 250% and 50 ms maximum injection time. The top 10 precursors were fragmented via HCD at 25% NCE.

For standard FTMS2 analysis, fragments were analyzed in the Orbitrap at 60,000 resolution, 1.2 m/z quadrupole isolation window, and a maximum injection time of 250 ms or 500 ms, as indicated in the respective figures.

For ITMS2-MS3 methods, fragments were analyzed in the ion trap with 1.5 m/z isolation window, a ‘Rapid’ scan rate, and a 150% normalized AGC target with the maximum injection time set to ‘Auto’. MS3 analysis was performed by selecting eight fragment ions using Synchronous Precursor Selection (SPS) with multi-notch isolation waveforms (1.5 m/z isolation window) to reduce interference in TMT ions due to potential co-isolation. These ions were further fragmented with HCD at 30% NCE and analyzed in the Orbitrap at 60,000 resolution, with 600% normalized AGC target and 250 ms maximum injection time.

For FTMS2-MS3 methods, fragments were analyzed in the Orbitrap at 15,000 resolution, 1.2 m/z isolation window, with the maximum injection time set to “Auto”. Subsequently, the eight most intense MS2 fragment ions were selected for SPS-MS3 scans using the previously mentioned parameters.

### MIRA-MS (Model-Informed Real-time Acquisition for Mass Spectrometry) implementation within inSeqAPI

MIRA-MS methods execute artificial intelligence prediction algorithms in real time within inSeqAPI, a program written in C# (.NET Framework 4.6.2) that interfaces with Thermo Orbitrap Eclipse and Ascend mass spectrometers (Tune version 4.3) through the instrument application programming interface (iAPI). Operation of inSeqAPI was maintained as previously described^18,22^, but with the addition of new filters including: PredictSpectraFilter, PredictRTFilter, and FDRFilter. During an LC-MS/MS analysis, MS2 spectra from the instrument were received by inSeqAPI and searched with either MSFragger-RTS which was operating on a remote server or locally with Comet. The resulting candidate peptides for each spectra were then considered for fragmentation and retention-time prediction. To calculate ‘spectral similarity’ in real-time, the predicted intensity array for each peptide was produced by Salud and compared to an array of experimentally matching peaks from the identification scan using a cosine similarity score. To calculate the ‘delta RT’ value in real-time, Salud was used to predict the iRT value for each peptide and this was compared to a real-time iRT value that was calculated during the LC-MS/MS analysis (**Supplementary Figure 2b**). To calculate the real-time iRT value, the predicted iRT values for non-decoy peptide candidates with spectral similarity >= 0.6 were added to a rolling list (**Supplementary Figure 2b**). The median of the last 50 iRT values was used to determine the real-time iRT value (**Supplementary Figure 2b**). New predictions were compared to this real-time iRT to produce a real-time iRT difference. To correct for potential lagging of the real-time iRT, the differences between the real-time iRT and predicted iRT were also added to a list and the median value was used to correct the real-time iRT difference, resulting in the final ‘delta RT’ value. The ‘spectral similarity’ and ‘delta RT’ were then considered within the FDRFilter which used a support vector machine to train a classifier to separate target and decoy peptides every 2,000 PSMs. Within the FDRFilter, multiple peptide matches (TopN) were ‘rescored’ per spectrum using the score provided by the SVM classifier to re-rank peptide matches.

### Optimization of MIRA-MS intelligent data acquisition methods

We evaluated different data acquisition architectures to identify the optimal method for immunopeptidomics using MIRA-MS (**Supplementary Table 3**).

While MS1 settings remained constant, MS2 scans were acquired either in the ion trap (low-resolution search parameters) or the Orbitrap (high resolution search parameters) and analyzed in real time. Scans were processed with the “MSFraggerFilter” and were followed by ppm error correction with the “PPMFilter”. Candidates were subsequently evaluated through the “PredictSpectraFilter” (Salud Nontryptic TMT Fragmentation Model, HCD 30 NCE, fragment ion mass tolerance 0.5 Da), and the “PredictRTFilter” (Salud Nontryptic TMT iRT Model, dynamic retention time alignment based on the previous 50 PSMs, with a spectral similarity threshold of 0.6). Peptide targets were validated using an “FDRFilter” (2000-scan training threshold, 200-scan forward count threshold), followed by a “DecoyFilter”. Peptides that passed these criteria triggered an MIRA-MS-mediated FTMS2 scan (Orbitrap; 60,000 resolution; 1.2 m/z isolation width; 30 NCE HCD). Variable injection times (50 ms, 100 ms, 250 ms, 750 ms, AutoMax) were tested based on the experiment.

For methods involving MIRA-MS-driven SPS-MS3, additional filtering was applied: a “Mass Range Filter” (455-2000 m/z), a “Precursor Range Exclusion Filter” (-15 m/z below to 5 m/z above precursor), a “Tag Loss Exclusion Filter” (TMTpro), an “SPS Filter” (quant modification = ‘TMTPro’, tolerance = 0.35 Da) and a “Top N Filter” (Top 8 peaks selected for next scan). Successful candidates triggered an SPS-MS3 scan (Orbitrap; 60,000 resolution; HCD at NCE 25 then NCE 45; 1.5 then 3 m/z isolation width; 250 ms max IT).

Real-time validation thresholds within the “FDRFilter” were systematically optimized for a range of values: 5%, 10%, 20%, 30%, 40% and 50%. Based on these iterations, thresholds of 10% and 20% were selected for FTMS2 and ITMS2 scans, respectively, to maximize target triggering while maintaining acceptable identification confidence. These parameters were applied consistently across all MIRA-MS workflows.

To maximize the identification of low-abundance peptides derived from non-canonical Open Reading Frames (nuORFs), we implemented a “ProteinPriorityFilter” within MIRA-MS. Peptides matching a nuORF database entry bypassed FDR filtering to directly trigger high-resolution FTMS2 scans with extended injection times. This strategy prioritized low-signal nuORF peptides for increased injection times to achieve the sensitivity required for confident identification and quantitation.

Furthermore, a “MaxITFilter” was implemented to allow dynamic adjustment of the injection time of the MIRA-MS triggered high-resolution FTMS2 scan based on the signal intensity observed in the initial instrument MS2 (either ITMS2 or FTMS2). This filter facilitated S/N-driven ion accumulation by defining a target signal-to-noise ratio of 1000 and a minimum S/N threshold of 10, allowing the instrument to dynamically extend injection times between 100 ms and 750 ms. Methods that employ the “MaxITFilter” are designated as AutoMax in the respective figures.

### MIRA-MS Analysis of Elastase Dilution Series

Method performance was evaluated across a range of sample loads (500 ng, 100 ng, 10 ng, 1 ng). For MIRA-MS guided FTMS2-FTMS2 acquisition, instrument MS2 injection times were compared at fixed intervals (50 ms, 100 ms, 200 ms) against the AutoMax setting, which dynamically optimized injection times via the “MaxITFilter”. For instrument FTMS2 methods, we tested 250 ms and 500 ms injection times and otherwise used the same parameters mentioned previously.

### Data Analysis

#### Online Data Analysis Parameters

Real time search was performed with MSFragger with the following parameters: Swissprot human database January 2023 version comprising 366,909 entries of canonical and protein isoforms, plus common contaminants, concatenated with a database containing NuORFs and SMORFs^9,23^. Decoy generation was performed on the entire database by adding reverse protein sequences for each entry. Overall, static modifications included Cys carbamidomethylation (+57.0215), K and n-term TMTPro (+304.2071); variable modifications included Met oxidation (+15.9949). The search enzyme name was set to unspecific, the length distribution was 7 - 15 amino acids, and top 2 results per spectrum were returned. FTMS2 spectra were searched with a precursor mass tolerance of 10 ppm and fragment mass tolerance of 20 ppm. ITMS2 spectra were searched with a precursor mass tolerance of 10 ppm and a fragment mass tolerance of 0.8 Da.

### Offline Data Analysis Parameters

Offline search parameters were optimized to maximize the number of identified and quantified peptides using the same parameters and database as online searches. For MIRA-MS-guided ITMS2-FTMS2 and FTMS2-FTMS2 architectures, both instrument- and MIRA-MS-triggered scans could potentially yield identifications, while quantification was consistently derived from the second (MIRA-MS-triggered FTMS2) scan. We therefore evaluated four search strategies: (i) high-resolution search (20 ppm fragment tolerance) of instrument-triggered scans only; (ii) high-resolution search of both instrument- and MIRA-MS-triggered scans; (iii) low-resolution search (0.8 Da fragment tolerance) of instrument-triggered scans only; and (iv) low-resolution search of both instrument- and MIRA-MS-triggered scans (**Supplementary Figure 3e,f**). Based on these evaluations, we selected the following optimal parameters for both instrument methods and MIRA-MS-guided architectures: FTMS2-MS3 (option i), ITMS2-FTMS2 and FTMS2-FTMS2 (option ii), ITMS2-MS3 (option iii).

### Clear Cell Renal Cell Carcinoma Immunopeptidomics

#### Sample Preparation

Clear cell renal cell carcinoma (ccRCC) cell lines A-498, TUHR10TKB and VMRC-RCZ were cultured in RPMI-1640 or EMEM media, supplemented with 10% fetal bovine serum and 2mM L-Glutamine. Cells were treated, in triplicate, with either 5 µM Vorinostat or DMSO for 18 hr prior to harvesting.

A*02:01-specific antibody (clone BB7.2) was coupled to AssayMAP Protein A cartridges (PA-W 5 µL) (Agilent, G5496-60000) and crosslinked with 20 mM dimethyl pimelimidate dihydrochloride (DMP) in 100 mM sodium borate pH 9.0. Crosslinked cartridges were quenched with 200 mM ethanolamine, washed with water, and stored in tris buffered saline containing 0.025% sodium azide at 4°C. pan-HLA-I antibody (W6/32) coupled cartridges were prepared the same way.

ccRCC cells were lysed in buffer containing 1% CHAPS, 20 mM Tris pH 8.0, 150 mM NaCl, 2 mM MgCl2, 50 U/mL Benzonase (Millipore Sigma), 1 mM EDTA, 0.2 mM iodoacetamide, 0.2 mM PMSF and 1x HALT protease inhibitor cocktail (Thermo Fisher Scientific). Samples were incubated on ice for 30 minutes with vortexing every five minutes. Lysates were clarified by centrifugation at 20,000 xg two times.

A*02:01 HLA-I peptide complexes were isolated from clarified lysate corresponding to ten million cells per sample using an automated liquid handling platform AssayMAP Bravo (Agilent). Clarified lysate was first passed over Protein A cartridges without antibody and then loaded onto prepared Protein A-BB7.2 cartridges (described above). Cartridges were washed with 400 mM NaCl 20 mM Tris pH 8.0, and then with 20 mM Tris pH 8.0. A*02:01 peptides were eluted with 0.1M acetic acid 0.1% TFA. The flow-through was then enriched for remaining HLA-I peptide complexes from other alleles using W6/32-coupled protein A cartridges in the same manner.

Eluted A*02:01 and pan-HLA-I peptides were loaded onto C18, reduced with 5 mM TCEP, and alkylated with 40 mM iodoacetamide. After washing with 200 mM HEPES, peptides were labeled with 50 µg TMTpro 18-plex (Thermo Fisher Scientific), washed with 2% acetonitrile 0.1% formic acid, and eluted with 50% acetonitrile 0.1% FA. Peptides were dried by vacuum centrifugation.

TMTpro-labeled peptides were reconstituted in 2% acetonitrile 1% formic acid, pooled, and loaded onto C18 stage-tips. Peptides were washed first with 1% formic acid, then with 2% acetonitrile 1% formic acid, and sequentially eluted in 1% formic acid containing increasing concentrations (10%, 15% and 50%) of acetonitrile. Peptides were dried by vacuum centrifugation.

#### Data Acquisition

A*02:01 and pan-HLA-I peptides were analyzed on an Orbitrap Eclipse mass spectrometer using an 185-minute standard FTMS2 or optimized FTMS2-FTMS2 MIRA-MS method. Peptides were reconstituted in 4 µl solvent A (2% acetonitrile 0.1% formic acid) and one µl of each fraction was loaded onto an Aurora XT 25 cm × 75 μm, 1.7 μm C18 column (IonOpticks). Peptides were separated for 155 minutes with 4 - 30% solvent B (98% acetonitrile 0.1% formic acid) in solvent A, followed by 5 min of 30 - 50% solvent B.

We employed a MIRA-MS-guided FTMS2-FTMS2 method with instrument-triggered MS2 scans acquired at 60,000 resolution in the Orbitrap (150% normalized AGC target, 200 ms maximum injection time). MIRA-MS-triggered FTMS2 scans were acquired for peptides passing a 10% real-time FDR threshold or mapping to nuORF sequences, using dynamic injection times of 100–750 ms. Control FTMS2 acquisition was performed as described above using 500 ms max injection time for dependent scans.

#### Immunopeptidomics Data Analysis

Spectra were searched offline using inSeqAPI with the same parameters as the online search and filtered to 1% FDR using Percolator. SCPCompanion was used for isobaric quantitation. Only peptides with sum signal-to-noise (sum SN) greater than 100 were used for quantitative analysis. For peptides with multiple measurements, the measurement with the greatest sum SN was used. Peptide summarization, median normalization and statistical analysis were performed using msstatsTMT^24^. Peptide HLA-I allele binding predictions were performed using HLApollo^25^ and peptides were classified according to MHC prediction rank: 0-0.05 for strong binders, 0.05-2 for binders and >2 for non binders.

nuORF transcripts were grouped into the following categories^9^: 3’ downstream ORF (3’ dORF), 3’ overlapping downstream ORF (3’ odORF), 5’ upstream ORF (5’ uORF), 5’ overlapping upstream ORF (5’ ouORF), canonical, long noncoding RNA (lncRNA), out-of-frame, and pseudogene.

#### Transcript abundance in tumor vs normal tissue (GTEX/TCGA)

NuORF transcript identifications were lifted over from hg19 to hg38 annotations using BioMart and used for filtering TCGA and GTEX transcript abundance data (downloaded from Xena^26^). TCGA data was further filtered for KIRC samples only. GTEX median abundance was calculated across all tissues excluding testis.

For Fig 2g, transcripts significantly changing with Vorinostat treatment (p < 0.05 in at least one cell line) were further filtered for a TCGA median log2 TPM >0 and TCGA median to GTEX median > 0.5. Significance in boxplot is based on Wilcox test.

Transcripts were designated tumor specific with the following criteria: 2x fold higher expression than median expression in GTEX except testis, present in at least 5% of TCGA samples, and GTEX max expression < 2 (log2 TPM+0.001).

#### Global Proteome Sample Preparation

For global proteome analysis, 50 µg of lysate was digested using suspension-trapping (S-Trap, Protifi) protocol. Sodium dodecyl sulfate was added to the lysate to 5% final concentration. Proteins were reduced with 5 mM dithiothreitol and alkylated with 10 mM iodoacetamide. Lysate was acidified with phosphoric acid to 2.7% final. Six sample volumes of binding buffer (100 mM TEAB, pH 7.5 in 90% methanol) were added and samples were loaded and bound to S-Trap. Bound proteins were washed five times with binding buffer and then digested on filter with Trypsin and LysC (Wako) at 1:25 enzyme:protein ratio for one hour at 47°C. Peptides were eluted sequentially with 50 mM TEAB pH 8, 0.2% formic acid, and 50% acetonitrile 0.2% formic acid. Eluted peptides were dried by vacuum centrifugation, reconstituted in 0.1% TFA and desalted with C18 cartridges on AssayMAP Bravo. Peptides were labeled with TMTpro, pooled and desalted with C18. Peptides were separated using Pierce high pH reversed-phase peptide fractionation kit into 24 fractions by eluting with increasing acetonitrile concentration from 5 to 35% and then with 40%, 50% and 60% acetonitrile. The 24 elutions were combined into 12 fractions and desalted with C18 cartridges on AssayMAP Bravo.

#### Global Proteome Data Acquisition

Global proteome fractions were analyzed on an Orbitrap Eclipse mass spectrometer equipped with FAIMS alternating between -40 and -60 CV using an 185-minute SPS-MS3 MIRA-MS method. All fractions were analyzed twice, first using tryptic parameters for RTS and second using semi-tryptic parameters. 500 ng of each fraction was separated on an Aurora 25 cm × 75 μm, 1.6 μm C18 column (IonOpticks) for 138 minutes with 4 - 30% solvent B in solvent A, followed by 15 min of 30 - 75% solvent B. MS1 scans were acquired in the Orbitrap with 120,000 resolution, 350 – 1350 m/z scan range, and 250% normalized AGC target or 50 ms max injection time. Data dependent MS2 scans were acquired for charge states 2 - 6 within a precursor fit window of 0.5 and a fit error of 70% in the ion trap using CID activation with 30% NCE, and 150% normalized AGC target or 100 ms maximum injection time. Spectra were analyzed in real-time using MIRA-MS. RTS parameters specified strict trypsin as the enzyme, up to 2 missed cleavages, peptide length of 7 - 30 amino acids, fixed TMTpro modification of the N-terminus and lysine, fixed carbamidomethylation of cysteine and variable oxidation of methionine (max 2). The number of enzyme termini was set to 2 for tryptic searches and 1 for semi-tryptic searches. Online FDR was set to 10% and no protein priority filter was used. Filtering was performed as described above with the addition of a protein closeout filter. Once 3 distinct and 10 total psms with 1% FDR were acquired for a protein, no further SPS-MS3 scans were triggered for that protein across all runs. For MS3 scans, eight MS2 ions were fragmented by HCD with 40% NCE and analyzed in the Orbitrap with 60,000 resolution, 120 - 2,000 m/z scan range, and 600% normalized AGC target or 400 ms maximum injection time.

#### Global Proteome Data Analysis

Raw files were converted to mzML using msConvert and searched offline with FragPipePlus version 23.2-build6. Offline search parameters matched online semi-tryptic search parameters with the addition of N-terminal acetylation as a variable modification. The top 2 matches were considered and fragmentation and retention time rescoring was performed with MSBooster using DIA-NN. Percolator was used for FDR calculation and ProteinProphet was used for protein grouping. PSMs and proteins were sequentially filtered to 1% FDR. Isobaric quantitation was performed for unique and razor peptides using Philosopher. Protein summation, median normalization and statistical analysis were done with msstatsTMT.

## Data and software availability statement

The original mass spectra, peptide reports and spectral libraries have been deposited in the public proteomics repository MassIVE (https://massive.ucsd.edu) with the associated MSV identifier MSV000101907 and can be accessed at. This dataset will be made public upon acceptance of the manuscript. As required by the instrument vendor, the inSeqAPI software enabling real-time inSeqAPI operation and MIRA-MS data collection is available upon request and requires users to have signed the Thermo IAPI agreement, as well as a distribution agreement with Genentech, Inc. We have deposited the source code for a matching version of inSeqAPI that can be operated in offline mode at github.com/Genentech/inSeqAPI_Offline and deposited this version at Zenodo (10.5281/zenodo.20348822). MSFragger-RTS database searching component has been made available as part of MSFragger software. The academic version is freely available at https://msfragger.nesvilab.org/, and the commercial users can obtain it from Fragmatics LLC at https://www.fragmatics.com/. A C# client communicating with MSFragger-RTS database searching is open source and can be found at https://github.com/Nesvilab/MSFragger-RTS-client.

## COI Statement

A.I.N. is the founder of Fragmatics LLC, which holds an exclusive commercial license from the University of Michigan for MSFragger. A.I.N. also serves on the scientific advisory boards of Protai Bio and Infinitopes. F.Y. is a paid consultant for Fragmatics LLC. A.I.N. and F.Y. have a financial interest related to the commercial licensing of MSFragger. A.M., K.L., S.K., and C.M.R. are employees of Genentech and shareholders of Roche.

## Figures

**Supplementary Figure 1:**
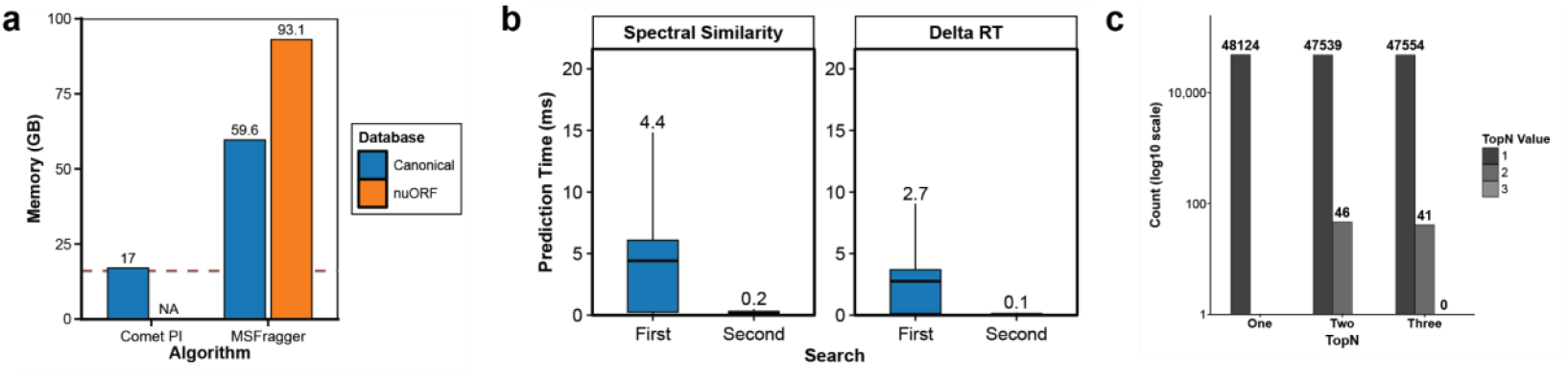
Characterization of search engine resource requirements, prediction performance, and top N settings. **a)** Memory requirements for real-time search database for Comet and MSFragger when searching a database of canonical sequences (canonical) or canonical + nuORF sequences (nuORF). **b)** Prediction times for the first search of a file (First) or when a file is searched a second time (Second) with the same exact parameters. Fragmentation and retention time predictions are stored in memory and can be recalled to decrease the time required for the prediction filter. **c)** The same files were searched considering one, two, or three peptides per peptide spectral match. The number of peptides assigned the rank of one, two, or three that passed FDR filtering is plotted for each search.

**Supplementary Figure 2:**
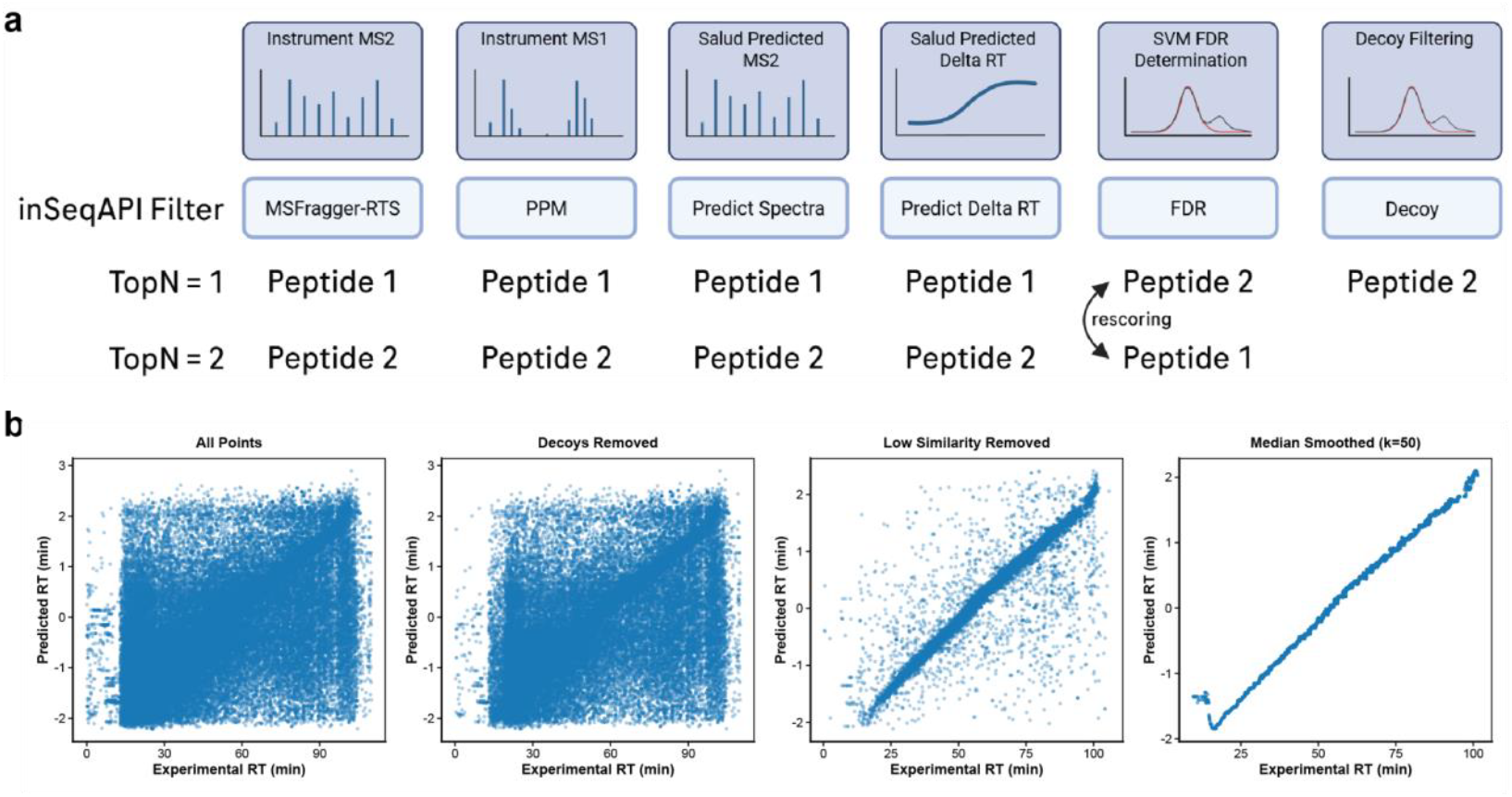
MIRA-MS workflow and real-time retention time difference calculation. **a)** Filters applied during an MIRA-MS workflow, including potential rescoring during the FDR filter. **b)** Filtering that enables real-time calculation of predicted and “actual” retention time. For all peptide spectral matches (PSMs) decoy PSMs are removed as well PSMs with a predicted spectral similarity < 0.6. Remaining PSMs are added to a list of “high-confidence” PSMs and a rolling median of the previous 50 measurements is used as the “actual” retention time. Because this value is a lagging indicator of true retention time, a correction factor is dynamically calculated and applied.

**Supplementary Figure 3:**
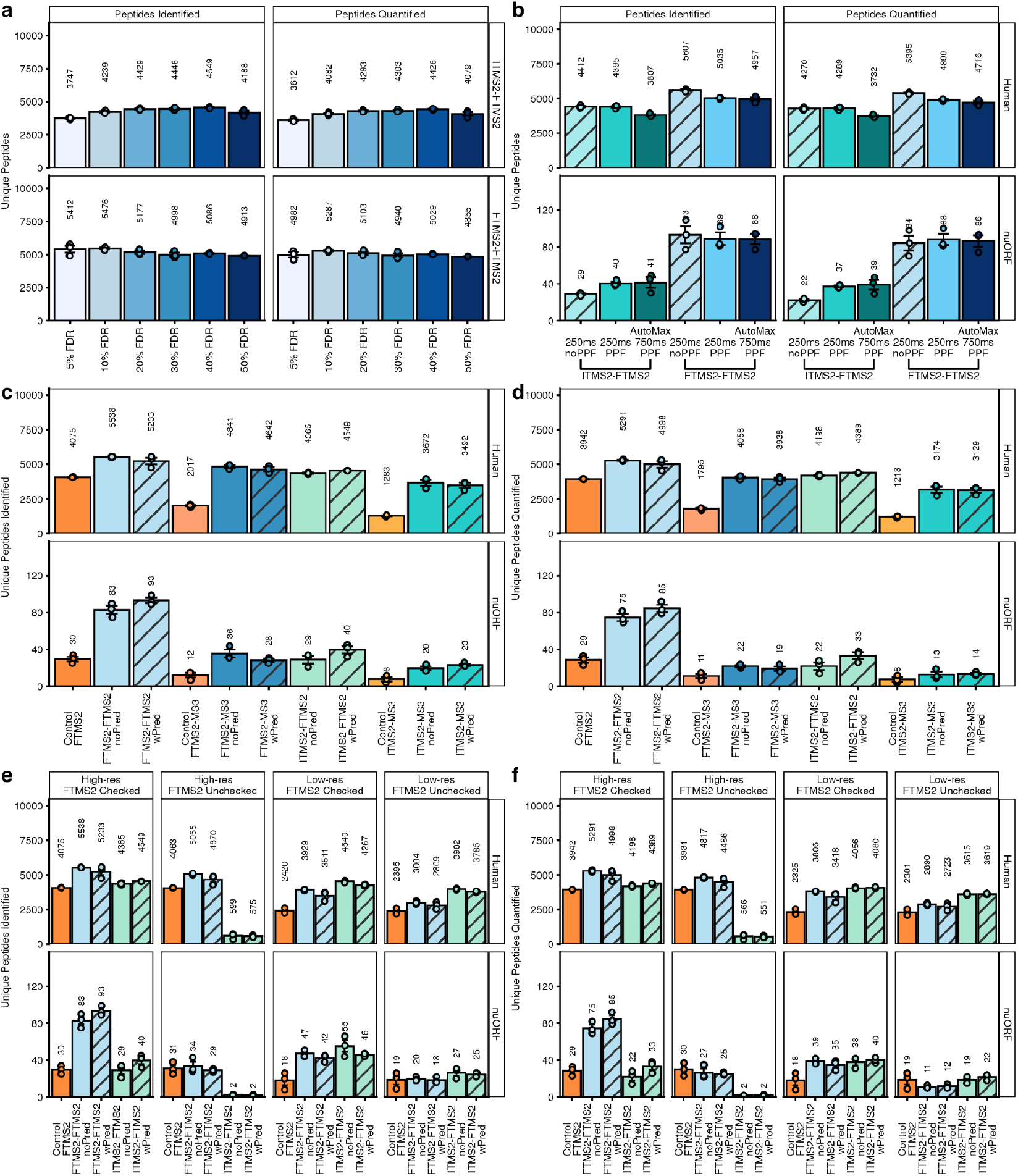
Optimization of MIRA-MS methods based on 500ng elastase digest. Comparison of total unique peptides identified and quantified (cumulative Sum S/N ≥ 100), categorized by canonical or nuORF origin. **a)** Selection of optimal online FDR thresholds. MIRA-MS performance was evaluated across various online FDRs thresholds, with 10% and 20% selected for FTMS2 and ITMS2 instrument scans, respectively, and applied to all subsequent experiments. **b)** Impact of the Protein Priority Filter on nuORF and canonical peptide recovery. The Protein Priority filter bypasses FDR for nuORF classified peptides. Performance comparison of ITMS2-FTMS2 and FTMS2-FTMS2 methods. Dynamic injection time was utilized to mitigate the time overhead of the Protein Priority Filter and was benchmarked against a fixed 250 ms injection time. **c)** Influence of real-time retention time and fragmentation prediction on peptide identification across method architectures. Comparisons were performed between instrument methods (FTMS2, FTMS2-MS3, ITMS2-MS3) and MIRA-MS-augmented configurations (ITMS2-FTMS2, FTMS2-FTMS2, ITMS2-MS3, FTMS2-MS3). Identifications are compared with (hashed bars) and without real-time prediction for both canonical and nuORF-derived peptides. **d)** Data are presented as in (c) for unique peptides meeting the quantification threshold (cumulative S/N ≥ 100). **e)** Influence of the search strategy on peptide identification across ITMS2-FTMS2 and FTMS2-FTMS2 method architectures. Identifications for canonical and nuORF-derived peptides were evaluated using four search strategies: (i) high-resolution search (20 ppm fragment tolerance) of instrument-triggered scans only; (ii) high-resolution search of both instrument- and MIRA-MS-triggered scans; (iii) low-resolution search (0.8 Da fragment tolerance) of instrument-triggered scans only; and (iv) low-resolution search of both instrument- and MIRA-MS-triggered scans. In all cases, MIRA-MS-triggered FTMS2 scans were utilized for quantification. **f)** Data are presented as in (e) for unique peptides meeting the quantification threshold (cumulative S/N ≥ 100). Based on these results, high-resolution search of both instrument and MIRA-MS-triggered scans (strategy ii) was selected for all subsequent experiments for ITMS2-FTMS2 and FTMS2-FTMS2 method architectures.

**Supplementary Figure 4:**
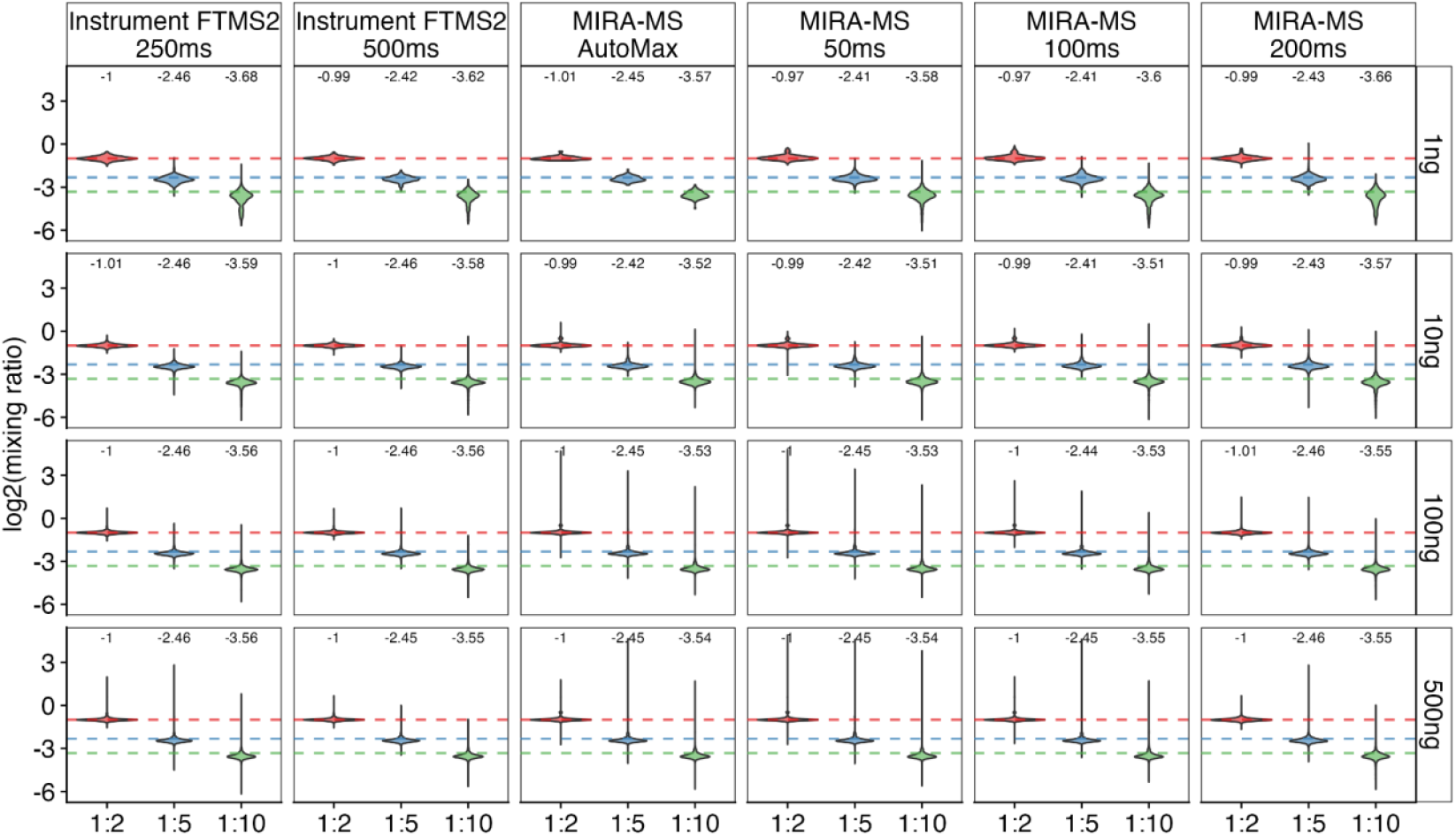
Expected vs. Observed distribution of TMTpro ratios across the serial dilution series of an elastase digest. Methods from Fig. 1i were included in this analysis, labels represent median values. Samples did not include interference proteome and thus data accuracy is reflective of measurement quality.

**Supplementary Figure 5:**
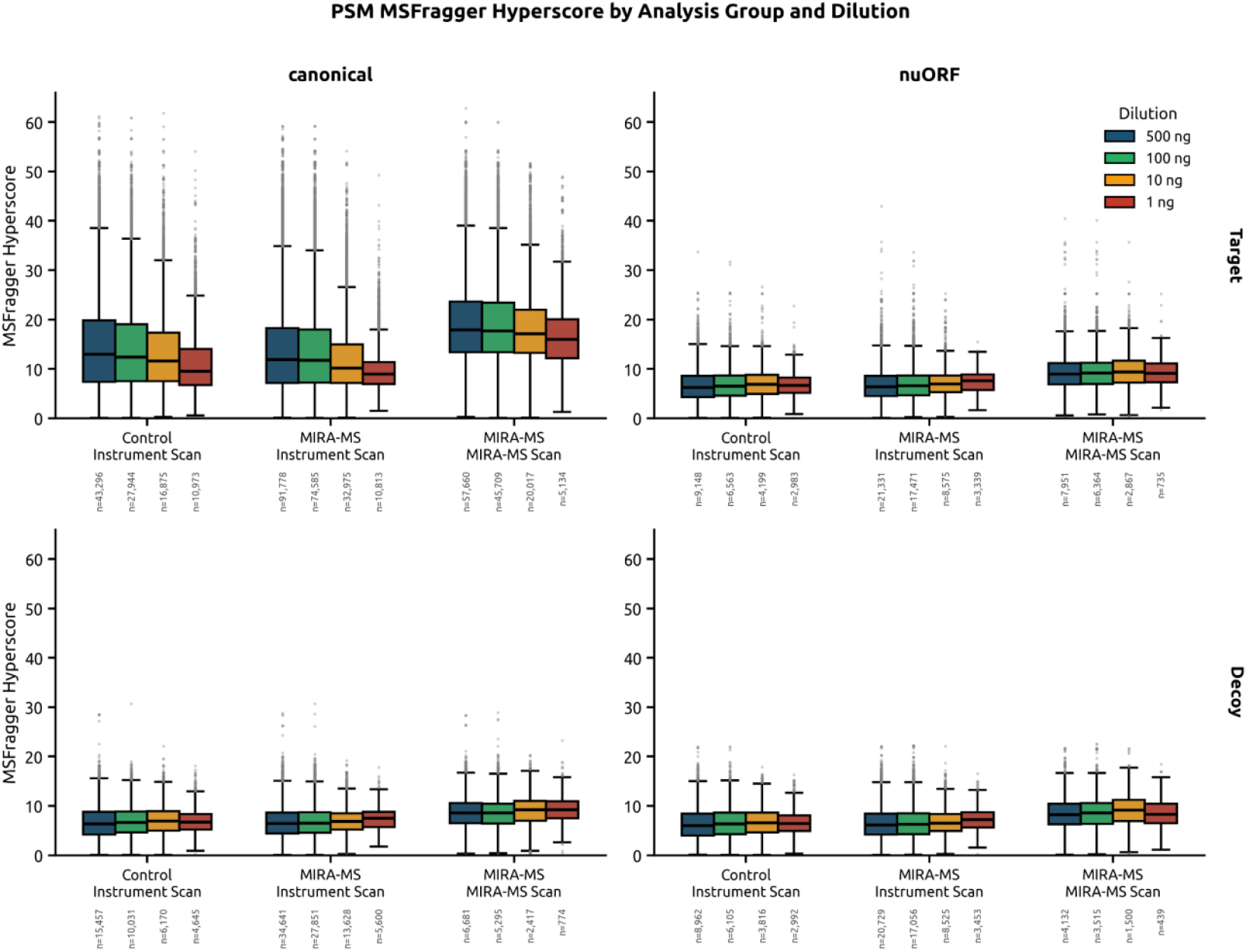
MSFragger hyperscores for all PSMs within dilution series of a nonspecific digest. PSMs were classified as Target (matching to a forward peptide) or Decoy (matching to a decoy peptide) as well as canonical (matching to a Uniprot protein) or nuORF (matching to a nuORF protein). Hyperscores were summarized by scan type (Control/Instrument Scan, MIRA-MS Instrument Scan (Identification Scan), MIRA-MS/MIRA-MS Scan (Quantification Scan), canonical vs. nuORF, and Target vs. Decoy.

**Supplementary Figure 6:**
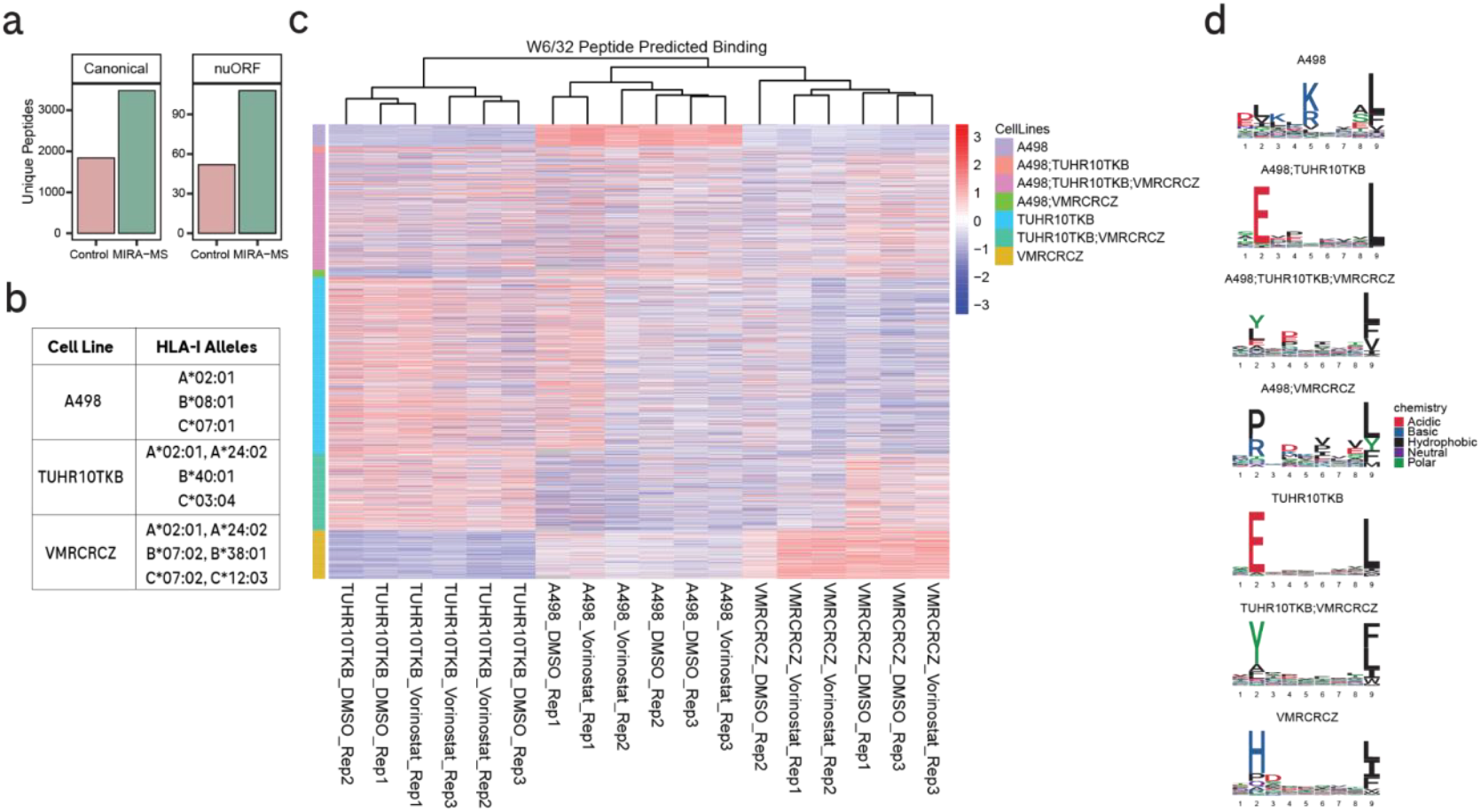
pan-HLA-I peptide sequence heterogeneity across cell lines. pan-HLA-I peptides were isolated from A*02:01 IP flow-through with W6/32 antibody, multiplexed with TMTpro, and analyzed using optimized MIRA-MS or standard FTMS2 analysis. (a) Number of unique quantitated peptides mapping to canonical or nuORF databases for each method. (b) HLA-I alleles expressed in each cell line. (c) Relative normalized abundance of pan-HLA-I peptides quantified with MIRA-MS, annotated by expected cell line based on HLApollo peptide-MHC binding prediction. Peptides identified in A*02:01 IP or without predicted binding allele were removed. (d) Sequence logos of 9-mer peptides for each group shown in (*c*).

**Supplementary Figure 7:**
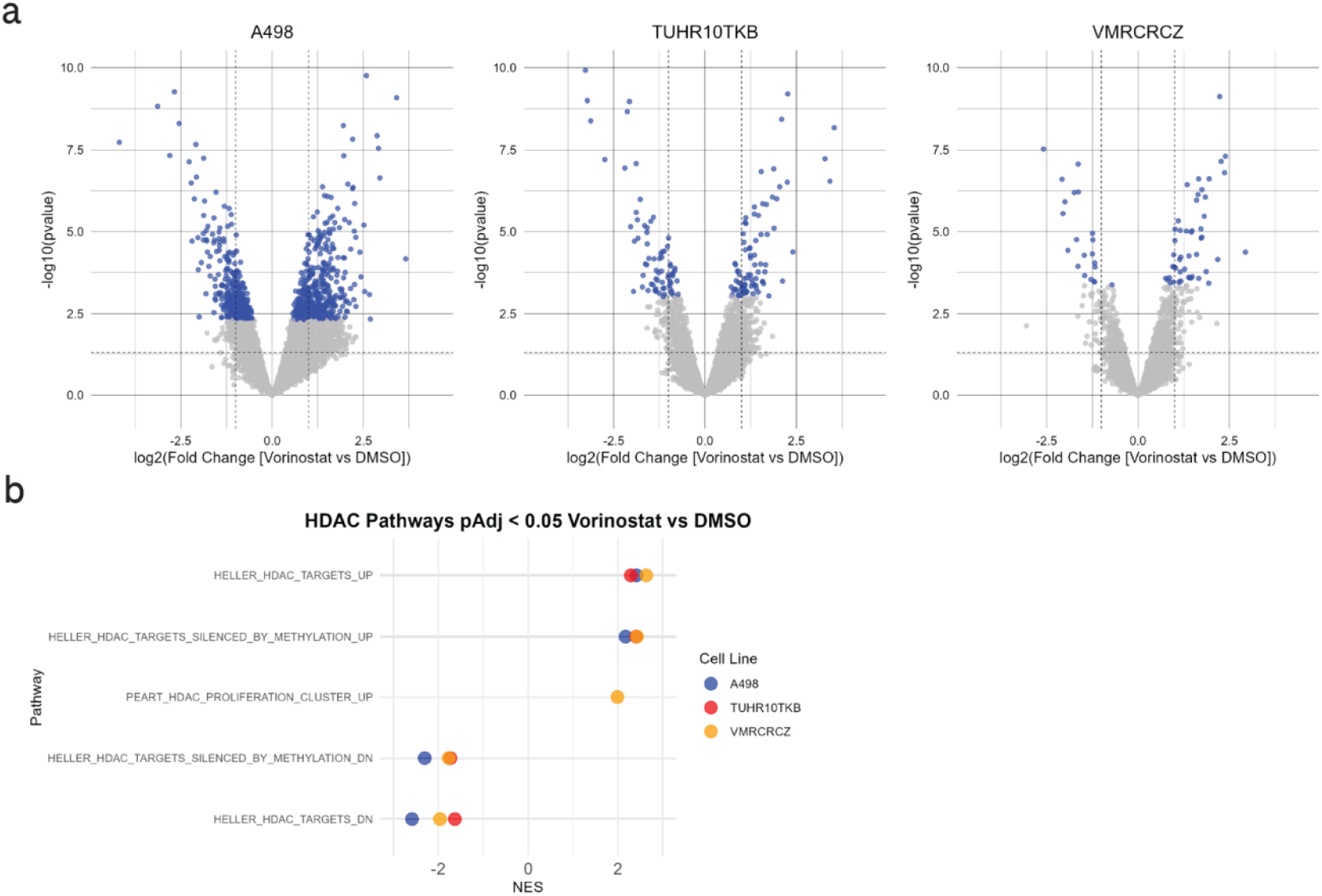
Vorinostat treatment alters global proteome abundance. (a) Differential protein abundance in Vorinostat vs DMSO treated cell lines. Significant proteins with pAdj < 0.05 are shown in blue. (b) HDAC signatures showing significant (pAdj < 0.05) enrichment by Gene Set Enrichment Analysis in the indicated cell lines with Vorinostat treatment. NES - normalized enrichment score.

